# Missing data approaches for longitudinal neuroimaging research: Examples from the Adolescent Brain and Cognitive Development (ABCD) Study

**DOI:** 10.1101/2024.06.12.598732

**Authors:** Lin Li, Mohammadreza Bayat, Timothy B. Hayes, Wesley K. Thompson, Arianna M. Gard, Anthony Steven Dick

## Abstract

This paper addresses the challenges of managing missing values within expansive longitudinal neuroimaging datasets, using the specific example of data derived from the Adolescent Brain and Cognitive Development (ABCD^®^) study. The conventional listwise deletion method, while widely used, is not recommended due to the risk that substantial bias can potentially be introduced with this method. Unfortunately, recommended alternative practices can be challenging to implement with large data sets. In this paper, we advocate for the adoption of more sophisticated statistical methodologies, including multiple imputation, propensity score weighting, and full information maximum likelihood (FIML). Through practical examples and code using (ABCD^®^) data, we illustrate some of the benefits and challenges of these methods, with a review of how these advanced methodologies bolster the robustness of analyses and contribute to the integrity of research findings in the field of developmental cognitive neuroscience.

## 1 INTRODUCTION

In many respects, it is an ideal time to be a developmental scientist. The current landscape offers substantial opportunities for in-depth study of developmental processes, owing to the proliferation of “big-data” computing resources (i.e., hardware and software) over the past two decades. This is further complemented by the emergence of accessible, expansive longitudinal datasets that span many years of development; in some cases, the entire human lifespan. Such datasets often contain information at multiple levels of analysis, including demographic, behavioral, contextual, and biological (e.g., neuroimaging), granting researchers unprecedented ability to explore inquiries that were once deemed implausible. However, these advances are not without their distinct analytical hurdles.

Working with extensive longitudinal datasets presents notable challenges, particularly when grappling with issues such as experimenter errors during data collection, participant nonresponse due to factors such as boredom, embarrassment, or fatigue (referred to as *item non-response*; [1]), and attrition over time. The term that encapsulates these difficulties is *missing data*. The Adolescent Brain and Cognitive Development (ABCD^®^) Study, initiated in 2015 and now in its ninth year of data collection, is one such dataset contending with missing data from multiple sources (i.e., item non-response from experimenter error, refusal to respond, or attrition).

Deciding on how best to accommodate or correct for missing data is one of the most daunting tasks for researchers. In this paper, our aim is to provide developmental scientists with practical tools and software examples tailored for the analysis of the ABCD^®^ Study. Although we will draw on existing papers on missing data in longitudinal research to offer best practices, our examples will primarily rely on open source software. These tools and recommendations are not only intended for users of the ABCD^®^ Study, but also for researchers working with other similar datasets (e.g., UKBB[2], PING[3], and HBCD[4] studies). We strive to improve the comfort and proficiency of researchers in analyzing large public data sets, along with the unique challenges such datasets pose.

### 1.1 Why should we deal with missing data in longitudinal studies?

Although missing data are common in research, the problem is compounded in longitudinal studies and creates challenges in interpreting longitudinal change. Unfortunately, to address missing data, researchers often rely on approaches that are statistically problematic. The most common is listwise deletion, which involves removing cases when any of the data fields are missing or removing cases with missing data only in the fields of interest, respectively [5, 6]. Researchers may do this and fail to realize the potentially serious implications for internal and external validity of the conclusions. Recall that internal validity refers to the degree to which a study accurately measures the cause-effect association it intends to investigate, without the influence of confounding variables or biases. In other words, a study has strong internal validity when changes in the dependent variable (the outcome being studied) can be confidently attributed to the manipulation of the independent variable (the factor being tested) and not to other extraneous factors. External validity refers to the extent to which the results of a study can be generalized beyond the specific conditions and participants of the study to a broader population, different settings, and/or different times. In other words, it assesses the degree to which the findings of a study hold for a wider range of individuals and circumstances.

Strong internal and external validity in psychological research is based on the notion of randomization. The assumptions of most of our statistical tests assume random and independent sampling from the population of interest. In the ABCD^®^ study, substantial effort was made to establish a population sample of adolescents in the United States using some degree of random sampling. The ABCD^®^ Study used a multi-stage probability sample of eligible children by probability sampling schools within the area of the area of catchment of each site [7]. Less than 10% of the sample was recruited through convenience methods (e.g., word-of-mouth). Thus, although the recruitment included multiple methods, random selection was a key feature of the ABCD^®^ Study and sets it apart from other large neuroimaging-focused studies of development [8]. The goal of this sampling strategy was to match the demographic profile of the ABCD^®^ sample with same-aged youth in two national surveys, the American Community Survey (ACS; a large-scale survey of approximately 3.5 million households conducted annually by the U.S. Census Bureau and collapsed into robust 5-year estimates) and annual enrollment data from third and fourth grade schools maintained by the National Center for Education Statistics [9]. The sampling strategy was additionally constrained by the requirement that study sites had available magnetic resonance imaging (MRI) scanners. Because these are typically available at research universities in urban areas, the sites were not chosen randomly and thus the sampling frame over-represents urban adolescents as opposed to rural adolescents and their families. Despite this caveat, the ABCD^®^ Study sample was largely successful at matching the ACS survey demographic profiles [10] (demographic assessments of the sample are summarized in Barch et al. [11]). That said, the ABCD^®^ Study sample is not completely representative of the target population of 9-10-year-old youth in the United States. For example, rural adolescents who did not live near an MRI scanner were not eligible for sampling. Additionally, the ABCD^®^ Study sample was significantly over-represented by very high-income households and youth with caregivers with professional degrees, compared to the target population [8]. Both of these examples introduce potential bias and concomitant threats to internal and external validity; the magnitude of bias can, in many cases, be difficult or impossible to ascertain.

In the same way, the common practice of removing participants from the sample due to missing data (i.e., by listwise or pair-wise deletion) introduces sampling bias. If missing data are related to the variables being studied, removing cases with missing values can lead to a sample that no longer accurately represents the target population being studied. Listwise and pair-wise deletion assume that missing data occur completely at random, meaning that the probability of data missingness is unrelated to the values of the observed and unobserved data. If this assumption is violated and missing data are related to the variables being studied, deletion can introduce bias and distort the true associations in the data. Furthermore, removing cases reduces the sample size, which impacts statistical power and increases the probability of a Type II error (that is, failure to detect a true effect). Finally, if the missing data are related to the variables being studied, deletion can alter the associations between variables. This can lead to inaccurate conclusions about the strength and direction of associations between variables. Listwise and pair-wise deletion are arguably the least desirable methods for dealing with missing data; the Task Force on Statistical Inference of the American Psychological Association called them ‘among the worst methods available for practical applications’ (p. 598)[12].

### 1.2 Mechanisms of missing data

This brings us to the obligatory and important discussion of the mechanisms of missing data (see also [13]). We use the traditional missing data terminology here [1, 14, 15], but anchor this discussion with specific examples from the ABCD^®^ Study for clarity. Three categories of missing data mechanisms are generally proposed: missing completely at random (MCAR), missing at random (MAR) and missing not at random (MNAR).

The best situation for missing data is when missingness occurs completely at random (MCAR). In this scenario, the probability of missing data is not related to the observed or unobserved variables in the statistical model. Any patterns of missingness are independent of the data, making it less likely that the missing data introduce bias or distort the associations within the dataset. In this case, even listwise or pair-wise deletion may not introduce substantial bias and, thus, MCAR is the least likely to compromise internal or external validity. As an example from the ABCD^®^ Study, consider that children are asked to complete an MRI scan at baseline, and 2-, 4-, and 6-year follow-up. However, due to unforeseen circumstances, some participants might not complete the scan, or certain parts of it, due to random factors not related to variables in the study (e.g., experimenter error, equipment failure, data transfer error, data corruption). The missing data in this case occur completely at random because there is no systematic pattern as to why certain scans are not acquired. Since there is no predictable pattern and the reasons for missing data are unrelated to other assessments or to the participants’ characteristics, this is an example of missing data occurring completely at random.

Although MCAR can occur, it is more common for data to be missing at random (MAR; also termed missing conditionally at random [16]). Suppose that an investigator uses the ABCD^®^ study to examine the association between household income and brain development. Participants with lower household income may be more likely to miss visits due to scheduling conflicts or transportation issues. In this scenario, the missingness mechanism (i.e., missing a visit due to transportation issues) is related to an observed variable (i.e., household income) but not directly to the outcome variable (e.g., volume of the caudate nucleus) that is missing. This situation represents MAR because the missingness depends on the observed variables present in the dataset. Another way to think about MAR is to consider whether the probability of missingness in the Y variable changes as a function of the X variable. The answer here is yes; as income goes down, the probability of missing MRI data goes up. This describes MAR because the missingness on Y is conditional on X, but it remains *equally probable* for all values of Y after conditioning on X. The missing Y values remain randomly distributed within each level of X.

If the data are MCAR or MAR, typical missing data approaches (e.g., maximum likelihood estimation or multiple imputation) work well because observed data distributions can be used as reasonable proxies for what the complete data would have been with no missing data [16]. But a third missing data mechanism can present problems. This mechanism, MNAR, would describe a situation in which the probability of missing data is unequal across the values of Y. Continuing with the above example, it might be the case that participants with lower household incomes might have different brain development patterns, *and* the missingness of their data is directly influenced by their income level. In this case, the probability of missing data is not random, but instead is associated with the variable of interest (e.g., caudate volume) in a non-random manner.

Although missing data mechanisms are readily identifiable in theory, in practice it can be difficult to ascertain which mechanism(s) are at play in real-world datasets. Students are commonly taught to apply a statistical test to examine the data. For example, groups with complete and missing cases can be examined for statistically different means or frequency distributions on some Y variable, or directly in a regression setting (e.g.,[17]). Researchers may be taught to apply Little’s [18] test to “confirm” data are MCAR. Little’s test sets the null to suggest the data are MCAR, so failure to find significance is commonly taken to indicate the researcher is “in the clear.” However, in practice, Little’s test is rarely useful. Enders (2022) argues such a test does not isolate potential auxiliary variables associated with missingness (i.e., it doesn’t say *which* variables are MCAR), and indeed only tests an omnibus hypothesis that is unlikely to hold in the first place [19]. Furthermore, Enders (2022) notes that simulation studies suggest that the test is underpowered, and particularly so for small numbers of variables, variables that have low correlation between the data and missingness, and when data are, in fact, MNAR. Although it is a common output for some statistical packages (e.g., SPSS), it may have a high probability of Type II error, ultimately providing statistical support for an assumption that may not be warranted (i.e., that the data are MCAR).

### 1.3 How much is too much missing data?

There is no universally agreed-upon amount or proportion of missing data that defines a rigid threshold. What constitutes “too much” depends on the study design, the research questions, the nature of the data, the mechanism of missing data, and the methods available for handling missing data. General guidelines roughly introduce cut-off points as “small and inconsequential amounts ≤ 5%”[20], “mild to moderate amounts 5% to 20%” [21], and “large and influential amounts (≥ 20%)”. Crucially, researchers should be transparent about the extent of missing data in their study by including an exploration of the missing data mechanism, explaining the approaches used for handling missing data, and analyzing the potential impact of different approaches on the results. Sterne et al. (2009) offered reporting guidelines for any analysis potentially affected by missing data. Researchers tend to be squeamish about analyses in which there are large amounts of missing data (say, ≥ 50%)[22], but it is precisely in these instances when researchers *should* be dealing with it, and it is not inherently inappropriate to tackle a data set with large amounts of missing data.

### 1.4 Missing data within dependent versus independent variables

Depending on the model of interest, data may be missing for the predictor/independent variable (X) or the outcome/dependent variable (Y). In general, missing data methods are designed to deal with missingness on either predictor or outcomes variables, or both. If data are missing only on the Y variable, it is easier to use full information maximum likelihood (FIML) approaches when this option is available [23]. For multiple imputation, missingness on the Y variable is somewhat more complicated. von Hippel (2007) published a detailed treatment on this issue [24], which reviews situations in which reliable and unbiased regression estimation under missingness on Y can be helped or hindered by imputation. von Hippel (and Little [25]) argue that, in cases where X’s are complete and only Y’s are missing at random, incomplete cases on Y contribute no information to the regression, and there is no need for imputation (i.e., a FIML procedure would likely be preferred). But in cases where X’s are also missing, imputing X’s with information from complete Y cases is potentially beneficial. However, von Hippel (2007) also argued that this is only helpful during the imputation step; he elaborated a step-wise method for incorporating the removal of the imputated Y’s (but not imputated X’s) in the ultimate analysis [24]. We do not advocate for such an approach here, principally because the benefits of this approach are situation-specific. Namely, they are applicable when the number of impu-tated datasets is small, when missing data in Y are MAR conditional only on the X variables, and when the proportion of missing values is large. Advances in computing power to improve the number of imputations and simultaneous advances in multiple imputation approaches render the extra removal step largely superfluous. Thus, the methods we review below are appropriate for situations in which data is missing in X, Y, or both.

### 1.5 Approaches to addressing missing data: Application to longitudinal neuroimaging studies

Longitudinal neuroimaging studies must attend to multiple sources of missing data. Imaging data may be missing at each wave of data collection due to high motion/low data quality, participant discomfort in the scanner that results in an incomplete scan, or cases where a participant does not attempt a scan for unknown reasons [8, 13]. The rates of missing neuroimaging data tend to be quite large, often larger than in missing behavioral data. In addition, imaging data may be missing across waves/time points due to classical attrition processes. With the advent of biosocial protocols in large survey datasets, some research suggests that collecting biological data (e.g., neuroimaging, genetics) may magnify attrition [26]. In addition, these issues are likely to compound over time, so long-term longitudinal studies like ABCD^®^ are likely to suffer acutely from missing data.

Neuroimaging data can be analyzed voxel-by-voxel [27, 28] with specialized software such as Statistical Parametric Mapping (SPM) in MATLAB [29], AFNI [30], or FSL [31]. Voxel-wise data can also be aggregated within a priori regions of interest (ROI) and analyzed as extracted vectors of data. Most users of large studies with imaging data analyze ROI-level data [32, 33]. In part, this is due to familiarity with standard statistical programs (e.g., R Statistical Software [34], SPSS [35], MPlus [36]). In addition, most specialized neuroimaging software does not allow for nested data structures (e.g., participants nested within study study and/or families, as in ABCD^®^) or missing data. Two new advancements in statistical software, FEMA [37] and Neuropointillist [38], are tackling issues with nested data structures. However, missing data in voxel- or vertex-wise analysis is a difficult problem, and there is not a straightforward solution in currently-available software packages. Thus, as most ABCD^®^ users use extracted ROI-level data, this paper will focus on all of the same approaches to missing data that are applied to behavioral data, and with available software and code.

Before tackling specific missing data approaches, we want to emphasize two points. First, missing data approaches (e.g., multiple imputation) should not be viewed as “making up data”. Instead, sophisticated statistical techniques are used to estimate or infer missing values based on the observed data patterns and their correlations. Appropriate missing data approaches, which are applied given the probable missing data mechanism and other assumptions, can make better use of the available non-missing information to make inferences based on the entire sample. These techniques play a vital role in promoting statistical generalizability[23]. Second, although we would like to present a manuscript that advances one missing data solution over others, there is no one statistical procedure for all situations. The appropriate procedure will depend on the structure of the data, the research questions addressed and ultimately the statistical model employed. Thus, our goal here is to present a few options and to make recommendations about when a researcher should choose one option over others.

#### Multiple Imputation

One of the approaches advocated for in contemporary statistical analysis is multiple imputation [14, 22]. Multiple imputation differs from related approaches such as single imputation using mean, median, or regression-predicted values to replace missing data. In those cases, a single value is assumed to be an accurate estimate of the missing value. Methods that use single imputation steps are flawed because they provide no way to estimate the degree of error or reliability of the imputations and may, in fact, *introduce* bias where none was previously present [39]. In contrast, in the words of Rubin (1987; p. 15): “Multiple imputation retains the virtues of single imputation and corrects its major flaws.” That is, multiple imputation incorporates the error inherent in estimating missing data into the calculation of the standard error estimates in the analysis. Therefore, multiple imputation is one recommended approach to dealing with missing data, and simulation studies show that it is effective in recovering accurate estimates that would have been derived from full data sets [40, 41]. We now turn to the details of this approach.

Multiple imputation entails a first stage of “filling in” missing data with samples from an appropriate distribution over an iterative procedure that leads to a number of separate completed data sets (say, 20). The substantive analytic model can then be fitted to each of the data sets, leading to, in this example, 20 parameter estimates for the same analytic model. In the second stage, the results are combined across the data sets according to Rubin’s rules [42], which establish how to combine estimates (e.g., slopes and standard errors) from each imputed dataset into overall multiple-imputation estimates (e.g., slopes) and associated standard errors. It is this second step that allows for the accounting of error uncertainty or error due to the imputation procedure. Multiple imputation is most appropriate for data that are MCAR or MAR, but it can also be applied to data that are MNAR[42].

These are the broad strokes. More detail is needed to understand multiple imputation applicable to longitudinal data, which are often analyzed using multilevel models. For this stage, many multiple imputation procedures use Bayesian estimation and Markov Chain Monte Carlo (MCMC) algorithms [16]. This procedure involves iterating between updating parameter estimates conditional on filled-in data, and then updating those missing values on the new parameter estimates. If the model is simple and it is reasonable to assume multivariate normality of all variables, this can proceed in a relatively straightforward manner using a series of univariate regression models that assume normal residuals (for example, fully conditioned model specification (FCS) [43]). In this case, the procedure is a direct method and the analyst can generally proceed agnostic to the analytic model.

However, more complicated models require so-called fully-Bayesian (model-based) approaches and proceed with information from the structure of the analytic model [23, 44]. This approach applies to models that incorporate moderators (i.e., that model interaction effects), are multi-level models, or are multi-level models with cross-level interactions. Model-based approaches can also employ other distributional assumptions in the estimation step (e.g., normal, skewed continuous, binary [probit or logit link], ordinal [probit link], multicategorical nominal [logistic link], or count [negative binomial link] distributions)[45, 23]. Importantly, agnostic imputation models impute a set of variables in a generic manner under the assumption of joint multivariate normality without concern for which variables are intended to function as outcomes (*Y*’s) versus predictors (*X*’s) in a set of later analyses. In contrast, the imputation model in a model-based approach should exactly match the intended analysis model. This includes distinguishing the model outcome from the set of model predictors and explicitly specifying features such as interactions (product terms), nonlinear relationships (e.g., polynomial terms), and multilevel random slopes. The software we review in our example below, Best Linear IMPutation (BLIMP) [45] is flexible and can implement both agnostic and model-based approaches for single-level and multilevel analysis models.

Whether employing agnostic or model-based imputation, the “Stage 1” implementation proceeds in the manner described above, with parameters iteratively estimated and updated across many computational cycles. Within this step, consisting of many thousands of iterations, the software will save a number of imputed data sets (after a specified number of “burns”) for “Stage 2.” The recommended number of imputation sets is at least 20 [46], and improved computer hardware can allow this process to proceed relatively quickly (depending on the nature of the model and the data). Prior to saving and analyzing these 20 (or more) imputed datasets, it is essential to assess the imputation model’s convergence. One useful metric for doing so is the potential scale reduction factor (PSR) [47]. This measure captures the similarity of MCMC chains that are initiated from different random starting values in accurately mapping the parameter space. Conceptually, the PSR is a ratio of the total variance and the average within-chain variance. As the between-chain variation (capturing discrepancies across chains) shrinks to zero, the total variance shrinks toward the average within chain variance and the PSR ratio approaches 1. Models with PSR values < 1.05 are considered to have reached convergence. If the number of iterations is insufficient to meet this cutoff value, one increases the number of iterations until it is met. In some cases, the number of iterations becomes extraordinarily high and the model may not converge. In this case, the diagnostic indicates that the imputation procedure is unable to arrive at a consistent answer with respect to the imputed data. At that point, it may be necessary to re-evaluate the model. We encountered this problem in our example, which is common for complex longitudinal data sets with missing data [22], and we specify below how we dealt with it.

If the model successfully converges and the imputation data sets are saved, the analyst can move to Stage 2. BLIMP software is designed to work with a variety of software platforms for the Stage 2 imputation step. Each imputation is saved as part of a “stacked” data set in a tab delimited text file. In our example we provide code for implementing Stage 2 multilevel imputation in R [34]. However, imputed datasets can be easily imported into other popular software suites (e.g., SPSS[35], Mplus[36]). As noted above, in Stage 2, the analysis model is implemented separately for each imputation data set, and then the model parameters are pooled using Rubin’s rules. For the multiple imputation point estimate, this is simply the average of the point estimates from each iteration data set (*M*). Thus,

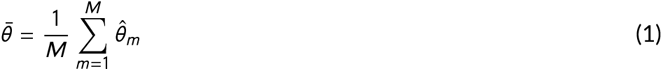

where 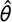 is the parameter value of the data set *m*, and 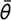 is the mean estimate of all the imputation data sets.

For the pooled standard error estimate, the equation incorporates corrections for the potential inflation of the estimate due to sampling error. Thus,

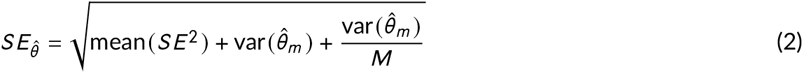

This essentially combines an average within-imputation sampling variance, a between-imputation sampling variance, and the squared standard error of the pooled estimate. This inflates the average standard error to compensate for sampling variability in the parameter estimates caused by missing data.

From this equation, one can derive the degree to which each predictor in a model is affected by missing data (i.e., by dividing the second and third terms under the radical by the quantity under the radical to yield a relative increase in variance due to non-response [42]). This fraction of missing information (FMI) can be calculated by hand, or obtained from the statistical package during estimation (e.g., R’s mitml package). The FMI values provide valuable information about the predictors in the model, but can also be used to inform the usefulness of auxiliary variables in the imputation step. In our analysis example, we implement multiple imputation using BLIMP for the first stage, and R for the second.

#### Full Information Maximum Likelihood (FIML)

Maximum Likelihood (ML) estimation can be distinguished from other statistical procedures commonly taught in introductory statistics courses (e.g., method of moments, method of least squares). These latter procedures stipulate that the estimates of the sample data (say p or 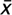) should be close to the parameters of the model (*π* or *µ*). ML procedures take essentially the opposite approach by attempting to establish which values of the model parameters would make the data most likely. Thus, ML procedures maximize the probability of the data given a model.

Linear models can be fit via least squares or ML to determine estimates 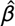 for the parameters *β*. However, least-squares estimation does not deal with missing data, instead using complete cases/case-wise deletion. Furthermore, only certain ML algorithms appropriately deal with missing data, a fact that is very confusing to most users of common statistical packages. For example, restricted maximum likelihood (REML) is a popular estimation procedure for multi-level models, and it is often an option to use instead of more general ML (e.g., in R’s lme4). But in both cases, REML and ML as implemented in many software packages handle missing data via complete case analysis–the missing data are deleted. In contrast, FIML is ML estimation with appropriate incorporation of cases with missing data [48, 49]. As FIML takes into account the information about missing data patterns, it can be particularly useful for longitudinal data with a hierarchical structure.

A full description of FIML is beyond the scope of this paper; more detail can be found in Hayes and Enders [16]. Briefly, likelihood can be thought of as a measure of fit between data and a set of model parameters. Whereas other procedures may fit the likelihood function starting with variance-covariance summaries, FIML applies likelihood estimation to each row in the data matrix. If data are missing from one row in the data matrix, the model estimation can still be used for the data that do exist. Thus, FIML neither discards nor imputes missing data. Instead, FIML typically applies an appropriate distribution matched to the data type to guess at the missing parts of the data as the algorithm iterates to a solution of the parameter values that have the best fit to the observed data. Enders and Bandalos [50] have shown that FIML produces unbiased parameter estimates under the MAR and MCAR mechanisms.

FIML is implemented in a number of subscription software packages (e.g., Mplus, SPSS AMOS, and for some procedures in SAS). However, it is not implemented in common open source packages for multilevel models (e.g., R’s lme4), and though it is an option in structural equation modeling packages through R (e.g., lavaan and OpenMx), as of v.0.6-17, lavaan does not currently handle complex multilevel models that researchers might employ for ABCD (e.g., nesting by study site). Thus, for FIML, we will use the popular Mplus package, which is widely used by the community for handling missing data, and is also the most straightforward. We will also provide Supplemental Code to conduct the same analysis in the open-source OpenMx package.

#### Propensity Scores (PS) Weighting

Lastly, PS estimation can be used to adjust for systematic differences between cases with complete and incomplete data, including observational studies in which data is MNAR. It involves estimating the probability of missingness conditional on complete covariates measured in all cases. In the context of a missing data analysis, “weights” refer to the numerical values assigned to observations in a dataset to reflect their probability of being included in the analysis. Estimated PS weights are then used to adjust the contribution of each “complete” or non-missing observations to the analysis. By including PS weights in the analysis, sociodemographic constructs or other measures of interest can be “balanced” between groups.

Although PS estimation was originally developed to estimate causal effects in observational data with treatment selection bias, at its core, PS can be used balance any two groups on a set of covariates (e.g., treatment/control; non-missing/missing). In the context of missing data, “treatment” is a binary variable, where “0” indicates cases with missing observations and “1” represents complete cases with regard to the response variable. There are several methods that differ in how they estimate the PS (e.g., logistic regression, Classification and Regression Trees [CART]), what they estimate (e.g., population treatment effect, treatment effect on the treated), and how the resulting PS are used in modeling (e.g., stratification, weighting).

Regardless of the method employed, one important step in PS estimation as applied to missing data is to identify a grouping variable that indicates whether cases are missing or complete. This grouping variable is typically the response variable that will be used in the ultimate statistical models; in the case of neuroimaging datasets, this is likely an indicator of missing neuroimaging data. Thus, PS estimates may differ depending on imaging modality (i.e., because there is far more missing data in functional modalities than structural modalities). Another critical early step in PS estimation is to decide which covariates will be used to balance groups. This is not a trivial task, and a one that is likely to impact the ultimate generalizability of study findings. For example, it is common to balance groups on a standard set of sociodemographic covariates (e.g., income, education, age, urbanicity, race-ethnicity). However, depending on the ultimate research aim (e.g., to examine group differences in neural synchrony between ADHD cases and controls), a researcher might also want to include other variables that demonstrate MNAR patterns (e.g., medication history; comorbid anxiety). Ultimately, balancing covariates should be informed by both theory and missing data analysis [51].

Once the missing data indicator and balancing covariates are identified, PS are estimated for every “treatment” case (i.e., non-missing indicator = 1). In situations where balancing covariates are also missing data, additional dummy variables indicating covariate missingness can be included as balancing covariates [52]. Groups are evaluated for balance by each covariate, weights are inverted (i.e., so that complete cases are weighted by the inverse of their probability of being a complete case), trimmed (i.e., to reduce the impact of outliers; very small or very large weights), and then used in statistical analysis [51]. An important note for PS approaches to handling missing data: the user’s ultimate analysis is only conducted on treatment cases, which are re-weighted to reflect (in our case here), the overall sample demographics. Thus, unlike other missing data approaches (e.g., FIML, MI), the effective sample size does not differ from a model that only includes complete cases.

### 1.6 Motivating Example

To illustrate the implementation and performance of missing data approaches in the ABCD^®^ study, we extend the results of a published study that examined microstructural development from 9 to 14 years of age. Using the first two waves of diffusion weighted imaging (DWI) from the ABCD^®^ study, Palmer and colleagues (2022) applied the restriction spectrum imaging (RSI) approach [53, 54] to test the associations between age and the proportion of restricted, hindered and free water diffusion within each voxel throughout the entire brain. The authors found widespread age-related changes in microstructural development. For example, the proportion of restricted diffusion within each voxel increased with age, with the greatest effects in the subcortical regions. As is common in neuroimaging studies, however, Palmer and colleagues implemented listwise deletion. This approach to missing data resulted in 8,086 participants (68% of the original sample) at baseline and 5,957 participants (50%) at the 2-year follow-up included in the study. This means that several thousand data points were dropped in this analysis, with no indication of the mechanism of missingness, or whether this introduced any bias to the analysis.

Our goal is to demonstrate how four different missing data approaches (i.e., listwise deletion, multilevel multiple imputation, propensity score matching, and FIML) could be implemented in longitudinal studies of age-related changes in brain development, using the ABCD^®^ Study data set as a specific example. As data loss varies significantly by modality [8], we include four imaging modalities from the ABCD^®^ Study in this exercise: diffusion-weighted imaging (DWI), task-based functional MRI (task-fMRI), resting state fMRI (rs-fMRI), and volumetric structural MRI (sMRI). We begin with an exploration of the missing data patterns and mechanisms. Second, we leverage each of the four missing data approaches to examine age-related associations with brain development. Four metrics of brain development were examined, one from each modality. Inspired by Palmer et al. (2022), we examined the restricted normalized signal fraction within the left caudate, which measures the degree of restricted diffusion averaged across all voxels within the ROI[53]. We also looked at three additional outcome variables previously shown to be associated with maturation, including reactivity of the amygdala to negative facial expressions versus neutral faces during the Emotional-N-back task (task-fMRI; [55]; temporal variance from resting state signal in the caudate nucleus (rs-fMRI), and cortical volume for the left *pars opercularis* of the inferior frontal gyrus (as defined by the Destrieux Freesurfer atlas [56]; sMRI). The names of each variable are provided in the Supplemental Code so that researchers can use these analyses as models for their own investigations. For each brain metric, we compared the point estimates that capture age-related changes from the models derived with each of the three missing data approaches.

## 2 METHODS

### 2.1 Participants and Procedure

The ABCD^®^ Study is a longitudinal population-based neuroimaging study of 11,865 9-10-year-olds in the United States [7]. The study employed a clustered probability sample to recruit eligible children from a comprehensive list of public and private schools within 21 catchment areas near study sites. Multiple children from each household were eligible to participate, including an oversample of twins from four of the 21 sites (150 - 250 twin pairs per site). The current study includes data from the Baseline and two-year-follow-up (2YRFU), downloaded from the ABCD^®^ Annual Curated Data Release 5.1 on the NIMH Data Archive (DOI: 10.15154/8873-zj65)[57]. There are 11,865 participants included in baseline, of which 10,902 were retained in the 2YRFU (note, the starting sample size at baseline is reduced slightly from earlier data releases because participants withdrew consent to participate and asked for their data to be removed).

**TABLE 1.** Demographics characteristics by timepoint.

|  | Response option(s) | Baseline | 2-year Follow up |
| --- | --- | --- | --- |
| n |  | 11866 | 10971 |
| Interview Age (Year) |  | 9.91 (0.62) | 12.03(0.67) |
| Sex (%) | Male | 6189 (52.2) | 5754 (52.4) |
|  | Female | 5677 (47.8) | 5217 (47.6) |
| Race (4-level, Baseline) (%) | White | 7516 (65.8) | 7068 (66.8) |
|  | Asian | 52 (0.5) | 43 (0.4) |
|  | Black | 1869 (16.4) | 1637 (15.5) |
|  | Other/Mixed | 1991 (17.4) | 1827 (17.3) |
| Ethnicity (Hispanic) (%) | Not hispanic | 9303 (79.4) | 8675 (80.0) |
|  | Hispanic | 2410 (20.6) | 2164 (20.0) |
| Family Income (%) | [>=100k] | 4559 (42.0) | 4832 (48.3) |
|  | [<50k] | 3222 (29.7) | 2487 (24.9) |
|  | [>=50k & <100k] | 3068 (28.3) | 2688 (26.9) |
| Parents Highest Education (%) | < HS Diploma | 786 (6.6) | 630 (6.9) |
|  | HS Diploma/GED | 1260 (10.6) | 1125 (12.2) |
|  | Bachelor | 3331 (28.1) | 3090 (33.6) |
|  | Post Graduate Degree | 2990 (25.2) | 2932 (31.9) |
|  | Some college | 3482 (29.4) | 1418 (15.4) |

Note. Means and standard deviations (parentheses) are reported for each variable.

### 2.2 Statistical Model

#### 2.2.1 Palmer’s analysis model

Several sociodemographic variables are recommended for inclusion in the analysis of ABCD^®^ data [58]. At the same time, many of these variables have some degree of missingness. Thus, in a typical analysis of ABCD^®^ data, the data will be missing for both the predictor/independent variable(s) and the outcome/dependent variable. For covariates, we started with the sociodemographic constructs listed in Table 2 of Palmer et al. [53], as these are recommended covariates and are common categorical operationalizations of questionnaire data administered as part of the ABCD demographics survey. We also note that other operationalizations have been implemented in the published literature (e.g., [59]). In fact, as we will see in our examples, in some cases the re-operationalization of measures is necessary to deal with missing data to either establish a complete data set up front, or to deal with problems with model conver-gence during analysis.

**TABLE 2.** Missingness Pattern by Demographics and Measure.

| Demographics | task-fMRI | rs-fMRI | dMRI | sMRI | Baseline | 2-Year Follow up |
| --- | --- | --- | --- | --- | --- | --- |
| Missing |  |  |  |  | 1498 | 2870 |
|  | Missing |  |  |  | 3941 | 4789 |
|  |  | Missing |  |  | 2250 | 4035 |
|  |  |  | Missing |  | 1470 | 3663 |
|  |  |  |  | Missing | 1138 | 3625 |
| Missing | Missing |  |  |  | 595 | 1366 |
| Missing |  | Missing |  |  | 321 | 1129 |
| Missing |  |  | Missing |  | 204 | 1041 |
| Missing |  |  |  | Missing | 159 | 1033 |
|  | Missing | Missing |  |  | 1665 | 3795 |
|  | Missing |  | Missing |  | 1133 | 3522 |
|  | Missing |  |  | Missing | 985 | 3489 |
|  |  | Missing | Missing |  | 1081 | 3499 |
|  |  | Missing |  | Missing | 931 | 3466 |
|  |  |  | Missing | Missing | 883 | 3429 |
| Missing | Missing | Missing |  |  | 260 | 1060 |
| Missing | Missing |  | Missing |  | 161 | 996 |
| Missing | Missing |  |  | Missing | 138 | 990 |
| Missing |  | Missing | Missing |  | 145 | 987 |
| Missing |  | Missing |  | Missing | 130 | 981 |
| Missing |  |  | Missing | Missing | 123 | 972 |
|  | Missing | Missing | Missing |  | 995 | 3473 |
|  | Missing | Missing |  | Missing | 876 | 3435 |
|  | Missing |  | Missing | Missing | 850 | 3412 |
|  |  | Missing | Missing | Missing | 824 | 3406 |
| Missing | Missing | Missing | Missing |  | 141 | 978 |
| Missing | Missing | Missing |  | Missing | 126 | 968 |
| Missing | Missing |  | Missing | Missing | 120 | 963 |
| Missing |  | Missing | Missing | Missing | 114 | 963 |
|  | Missing | Missing | Missing | Missing | 805 | 3398 |
| Missing | Missing | Missing | Missing | Missing | 113 | 958 |

Note. dMRI = Diffusion-weighted magnetic resonance imaging; task-fMRI = task-related functional magnetic resonance imaging; rs-fMRI = resting state magnetic resonance imaging; sMRI = structural magnetic resonance imaging.

The general model from Palmer et al. [53] is:

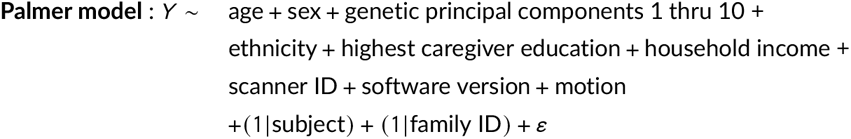

The independent variable that is the primary focus in this model is age (measured in months). Sociodemographics included sex of youth assigned at birth (male/female), highest caregiver education (< HS diploma, HS diploma/GED, Bachelor, Some College, Post Graduate Degree), household income (less than $50K, $50 - $100K, more than $100K), Hispanic ethnicity (Yes/No). The movement index specific to the outcome variable (e.g., average framewise displacement) is also included. Palmer and colleagues used the top 10 genetic principal components from the genetic assay [60] to indicate ancestry. We note that racial-ethnic identity is not equivalent to genetic ancestry [59]. Palmer also modeled scanner ID (32 levels) and software version (23 levels) as fixed-effects, and subject and family as random effects.

#### 2.2.2 Issues encountered and modified model for subsequent analysis

Proceeding with this model, we were able to replicate the findings with listwise deletion. However, we immediately encountered difficulties with model convergence during the multiple imputation procedure. These issues were caused by the fact that some of our categorical predictors could not be modeled together because the bivariate cross-tab tables of these predictors contained cells with cell counts that were either very low or actually zero. This provides insufficient information for the imputation modeling. Notably, these are the kinds of issues that investigators might regularly encounter, and either not know how to proceed, or perhaps they may even abandon attempting to deal with missing data altogether. We hope our illustration encourages investigators to search for other solutions, as we do here.

Because age is our primary predictor, we simplified the model with respect to how the covariates and random effects were defined and modeled. The first step was to simplify the nesting structure of the model to allow only subject id as the level-2 sampling unit. Nesting at a third level is possible in BLIMP, but in this example the more complex model with subject and family ID failed to converge. Thus, some variance associated with siblings is not modeled here because we removed the family variable. We still encountered issues. For example, there are categorical variables with a large number of levels (e.g., scanner ID and software version). This is not necessarily a problem except when combinations of categorical variables must be considered in order to understand the correlation structure of the data. For example, if we consider a cross-tab table for ethnicity (2 levels) and scanner ID (32 levels), there are a number of cells with zero cell counts in this cross-tab table (i.e., zero people of a certain ethnicity are represented at particular scanners). Thus, the data do not contain the necessary information to facilitate the calculation of the correlation between the variables, and this will lead to convergence issues. We encountered this issue in our example. The solution is either to collapse categories to eliminate cells with zero or very low cell counts, or to choose alternative variables to include in the model (i.e., to change the model). In lieu of scanner ID, to simplify the number of levels, we converted scanner ID to a 3-category “scanner manufacturer” variable (i.e., GE, Siemens, Philips), which reduced 32 levels to three. We also found it necessary to eliminate the software version variable. Thus, scanner and software variance is only partially captured by the scanner manufacturer variable entered as a fixed effect. These changes show the kinds of analytic decisions that go into modeling ABCD data, and indeed we can see how this impacts the estimation of the effects of interest (e.g., age) and how it impacts the implementation of various missing data approaches.

Through this process of testing models to identify successful convergence, we settled on a Modified Model, which we used to assess each of the four approaches:

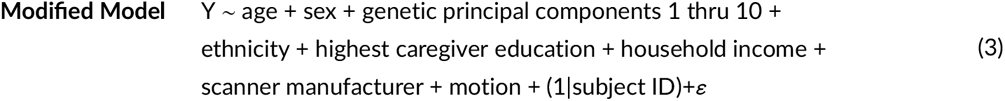

A second issue we encountered relates to data organization. For some software, reformatting of the data is necessary for longitudinal models. Some software is equipped to deal with long- and wide-formatted data, while other software packages work better with one or the other. Furthermore, an idiosyncratic aspect of ABCD is that some participants do not have any data for follow-up visits, and in the ABCD data release (which is in long format) these participants are coded as having missing data rows. For example, participant ‘XYZ’ may have a row for the baseline visit, and could have complete or missing data within the row, but have no data row for the 2YRFU (i.e., instead of having a row filled with NA values, the participant row for 2YRFU is absent altogether). This is not an issue for case-wise deletion, as the row would be eliminated anyway, but for certain missing data approaches, reformatting of the data is necessary. In some respects, this was the most challenging aspect of the analysis, and we include the code for reformatting in Supplemental Code.

At this point we can proceed with our examples. Thus, in the following section we compare three broad missing data approaches–multiple imputation, propensity score matching, and FIML–against listwise deletion to illustrate how one might approach longitudinal modeling within the ABCD data set.

### 2.3 Statistical Analyses

#### 2.3.1 Listwise deletion using the linear mixed-effects model

The first set of models implemented linear mixed effects modeling using the lmer() function from the lme4:: package [61] in R (v4.2.1 (2022-06-23)) to estimate associations between age and brain metrics. lme4 proceeds with complete-case analysis, otherwise known as listwise deletion. For each imaging modality, the extracted ROI (e.g., bilateral amygdala reactivity to negative versus neutral faces) was specified as the dependent variable, with age in years as the primary predictor variable, and the rest of the model structure as specified in the Modified Model.

#### 2.3.2 Multiple imputation using *BLIMP*

As noted above, multiple imputation is a two-stage procedure. Stage 1, building the multiply imputed data set, happens in BLIMP. We proceeded with a model-based approach–the imputation model is expected to match the analytic model. As we noted above, we began with the Palmer Model, and due to convergence issues settled on the Modified Model. The output of this successful model is reported below in Results.

#### 2.3.3 Propensity score matching and inverse propensity weighting

Propensity scores to adjust for missing data were estimated at both baseline and 2YRFU using the “Toolkit for Weighting and Analysis of Nonequivalent Groups (twang)” package (v2.5) [62] in R (v4.2.1) [34]. Specifically, the ps() function implements gradient-boosted models to balance any two groups on a set of covariates. ^1^. “Treatment” is a specified binary variable, where “0” indicates cases with missing observations and “1” represents complete cases with regard to the response variable (e.g., bilateral amygdala reactivity). The estimating method “ATE” (Average Treatment Effect) is used to compute weights for the treatment cases to estimate the population average treatment effect (i.e., we are interested in estimating models that generalize to the entire sample of both treatment and control cases, so the weights seek to balance the treatment group sociodemographic composition to the entire sample). The ‘ATE’ algorithm identifies the iteration (i.e., complexity) of the gradient boosted models that achieves the best balance between groups.

The covariates used to balance the control (i.e. missing) cases with the treatment (i.e. complete) cases fall into two categories. First, we specified a set of standard sociodemographic variables, all of which were used to construct post-stratification weights that match the entire baseline recruited sample with the target population of 9 to 10 years old in the American Community Survey [9, 63]. A useful feature of the twang package is that first-order interactions between all covariates are added to the model automatically (e.g., caregiver education X household income); thus, users do not need to specify every interaction term that might predict missingness. To account for missing data in both the neuroimaging response variables and covariates, we adopt an approach by D’Agostino and Rubin (2000) [52] by including in the set of balancing covariates a series of binary variables that indicate whether a covariate (e.g., household income) is missing. Including binary variables indicating covariate missingness ensures that all cases with complete data on the response variable (e.g., bilateral amygdala reactivity) are included in the analysis regardless of whether there is missing covariate data.

#### 2.3.4 Full information maximum likelihood (FIML) estimation

Lastly, we applied full information maximum likelihood (FIML) estimation as another approach to address missing data. We implemented FIML using the *Mplus*[36],which provides extended structural equation modeling (SEM) algorithms and is probably the most user-friendly option now to estimate model parameters using FIML. We implemented FIML estimation for the Modified Model as described.

## 3 RESULTS

### 3.1 Examination of Missingness Patterns

First, we examine missing data patterns in the four imaging modalities with respect to all sociodemographic covariates. Table 2 shows that there were 31 unique patterns of missing data. At both the baseline and 2YRFU the most common missing data pattern among participants was missingness for task-fMRI. For example, 3,941 and 4,789 participants were missing only task-fMRI data at baseline and 2YRFU, respectively. By comparison, the same missing data patterns for dMRI included 1,470 (baseline) and 3,663 (2YRFU) participants. Other common missing data patterns included missing both task-fMRI and rs-fMRI, as well as missing sociodemographic data in isolation.

Logistic regression models were used to examine predictors of missing neuroimaging data at each wave. For task-fMRI data at baseline, age (OR = 1.42, p < 0.001), sex (Female, OR = 1.14, p = 0.003; ref = Male or intersex), race (Asian, OR = 0.38, p = 0.001; Black, OR = 0.50, p < 0.001; Other/Mixed, OR = 0.87, p = 0.023; ref = White), annual household income (<50k, OR = 0.71, p < 0.001; ref = [>=100k]), and caregiver highest education (< HS Diploma, OR = 0.57, p < 0.001; HS Diploma/GED, OR = 0.66, p < 0.001; <Some college, OR = 0.76, p < 0.001; ref = Bachelor), were significantly associated with missingness. Missing task-fMRI data at the 2YRFU was associated with age (OR = 0.67p < 0.001), sex (Female, OR =0.91, p = 0.033; ref = Male or intersex), race (Black, OR = 0.51, p < 0.001 ; Other/Mixed, OR = 0.86, p = 0.018; ref = White), ethnicity (Hispanic, OR = 0.85, p = 0.015; ref = Non-hispanic), annual household income (≥ 50k&<100k, OR = 1.19, p = 0.004; ref = [>=100k]), and caregiver highest education (< HS Diploma, OR = 0.67, p < 0.001; HS Diploma/GED, OR = 0.69 p < 0.001; Post graduate degree, OR = 0.87, p = 0.013; ref = Bachelor). These analyses suggest that the missingness mechanism for task-fMRI is not MCAR. [64, 65].

For DWI, similar patterns of associations between sociodemographic constructs and missingness were revealed in both waves. For example, age (OR =1.24, p < 0.001), sex (Female, OR = 1.26, p < 0.001; ref = Male or intersex), race (Asian, OR = 0.42, p = 0.017; Black, OR = 0.70, p < 0.001 ; Other/Mixed, OR = 0.78, p = 0.002; ref = White), ethnicity (Hispanic, OR = 1.23, p = 0.016; ref = Non-Hispanic), annual household income (<50k, OR = 0.73, p = 0.001; ref = [>=100k]) were significantly associated with missing DWI data at baseline. In the 2YRFU, age (OR = 0.54, p < 0.001), sex (Female, OR = 0.83 p < 0.001; ref = Male or intersex), race (Black, OR = 0.66, p < 0.001 ; Other/Mixed, OR = 0.87, p = 0.042; ref = White), ethnicity (Hispanic, OR = 0.82, p = 0.005; ref = Non-hispanic), annual household income (≥ 50*k* &< 100k, OR = 1.24, p < 0.001; ref = [>=100k]), and caregiver highest education (HS Diploma/GED, OR = 0.81, p = 0.028; post graduate degree, OR = 0.84, p = 0.003; ref = Bachelor) were significantly associated with DWI data missingness. Thus, similar to task-fMRI, missing DWI at both waves are unlikely to be MCAR.

Overall, the patterns suggest that imaging at each of the first two timepoints is not missing completely at random (MCAR), and implementing listwise deletion may introduce bias. To mitigate bias and maintain internal and external validity, good statistical practice would suggest that including predictors of missingness in the models (e.g. FIML) or other appropriate missing data approaches as discussed above.

### 3.2 Evaluation of Multiple Imputation

Using the Modified Model, with the number of burns specified in our Supplemental Code, we obtained 20 imputed data sets which converged with a PSR of 1.031 (for dMRI), 1.032 (for task-fMRI), 1.018 (for rs-fMRI), and 1.040 (for sMRI). All PSRs are less than the suggested cutoff of 1.05 [47], indicating that the procedure was able to arrive at consistent answers across the MCMC chains. This licenses us to proceed to Stage 2, which we completed in R using mitml. BLIMP packages the output imputations of Stage 1 for easy use with this and other software packages, and we provide syntax in Supplemental Code. The Stage 2 analysis provides the point estimates and confidence intervals, shown in Table 3 and 4 for the dMRI analysis as an example. Note that the standard errors are computed according to Rubin’s rules as described, which is also applied to the calculation of the 95% Confidence Intervals. In addition, FMI values are provided for each predictor, indicating the degree to which each estimate is affected by missing data.

**TABLE 3.**
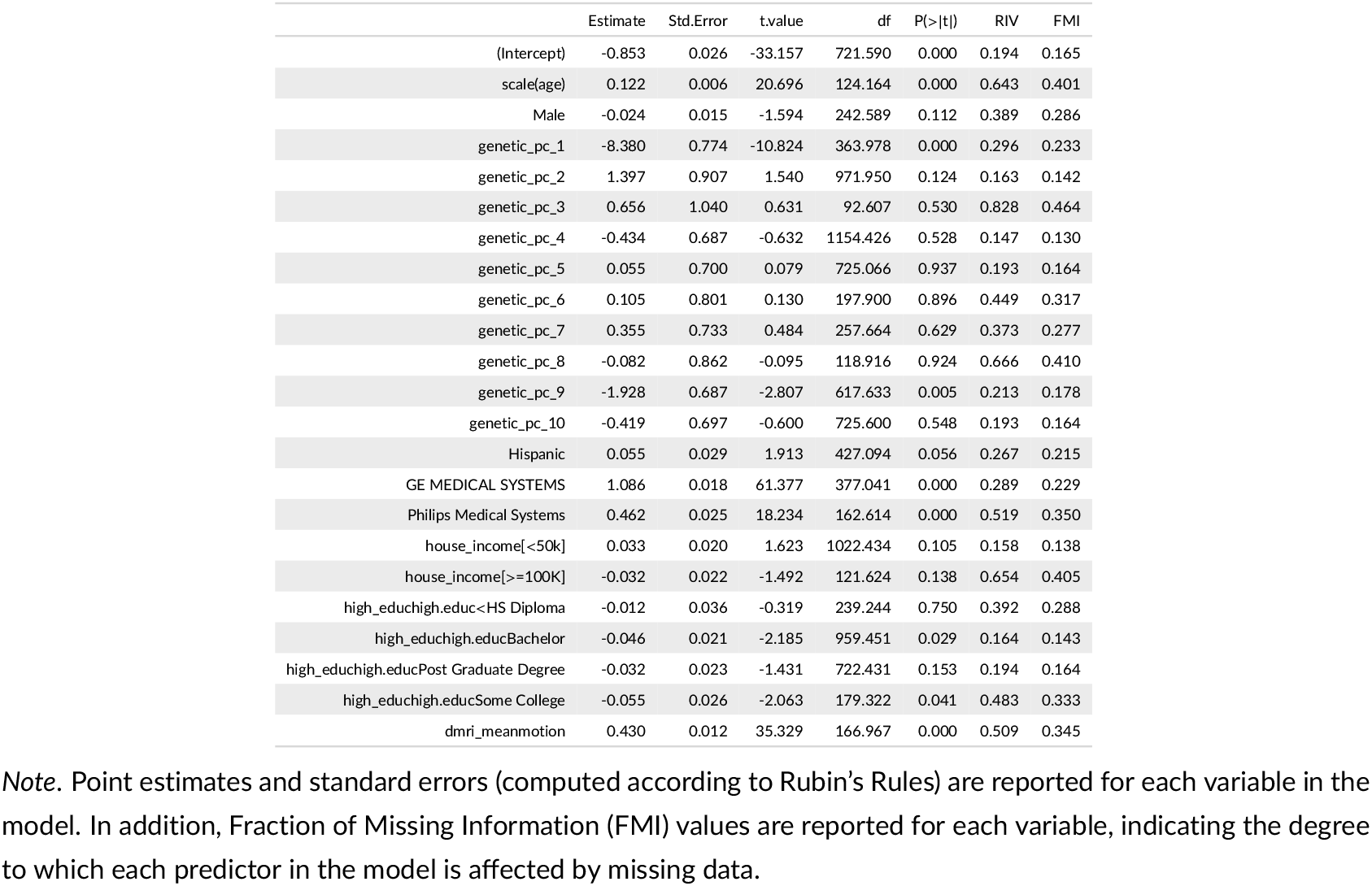
Example output from Stage 2 Multiple Imputation step from R package mitml for diffusion-weighted magnetic resonance imaging measure.

**TABLE 4.** Example output from Stage 2 Multiple Imputation step from R package mitml for diffusion-weighted magnetic resonance imaging measure: 95% Confidence Intervals for each Point Estimate.

|  | 2.5 % | 97.5 % |
| --- | --- | --- |
| (Intercept) | -0.90 | -0.80 |
| scale(age) | 0.11 | 0.13 |
| Male | -0.05 | 0.01 |
| genetic_pc_1 | -9.90 | -6.86 |
| genetic_pc_2 | -0.38 | 3.18 |
| genetic_pc_3 | -1.41 | 2.72 |
| genetic_pc_4 | -1.78 | 0.91 |
| genetic_pc_5 | -1.32 | 1.43 |
| genetic_pc_6 | -1.48 | 1.68 |
| genetic_pc_7 | -1.09 | 1.80 |
| genetic_pc_8 | -1.79 | 1.62 |
| genetic_pc_9 | -3.28 | -0.58 |
| genetic_pc_10 | -1.79 | 0.95 |
| Hispanic | -0.00 | 0.11 |
| GE MEDICAL SYSTEMS | 1.05 | 1.12 |
| Philips Medical Systems | 0.41 | 0.51 |
| house_income[<50k] | -0.01 | 0.07 |
| house_income[>=100K] | -0.07 | 0.01 |
| high_educhigh.educ<HS Diploma | -0.08 | 0.06 |
| high_educhigh.educBachelor | -0.09 | -0.00 |
| high_educhigh.educPost Graduate Degree | -0.08 | 0.01 |
| high_educhigh.educSome College | -0.11 | -0.00 |
| dmri_meanmotion | 0.41 | 0.45 |

### 3.3 Evaluation of the performance of the PS matching process

The propensity scores were estimated on both baseline and 2YRFU data for each of the four imaging modalities (i.e., eight models in total), using the twang::ps() function with gradient boosting algorithms [66]. Across all models, the number of iterations that minimized the mean absolute effect size (i.e., es.mean) between the treatment (i.e., complete) and control (i.e., missing) cases ranged from 917 iterations in the sMRI baseline model to 3,239 iterations in the task-fMRI YR2FU model (see Supplemental Tables). Figures 1 (a) - (d) depict the balance of covariates before and after applying propensity score matching to the dMRI and task-based fMRI metrics, while Figures 2 (a) - (d) demonstrate the balance plots for the rs-fmri and sMRI metrics. Across all models, the absolute value of the standardized mean differences between groups is < 0.10 at both the baseline and the 2YRFU, indicating successful covariate balance with the PS estimate. Overall, the plots suggest that age, race, and household income contributed the largest variance to the PS scores at both baseline and YR2FU.

**FIGURE 1.**
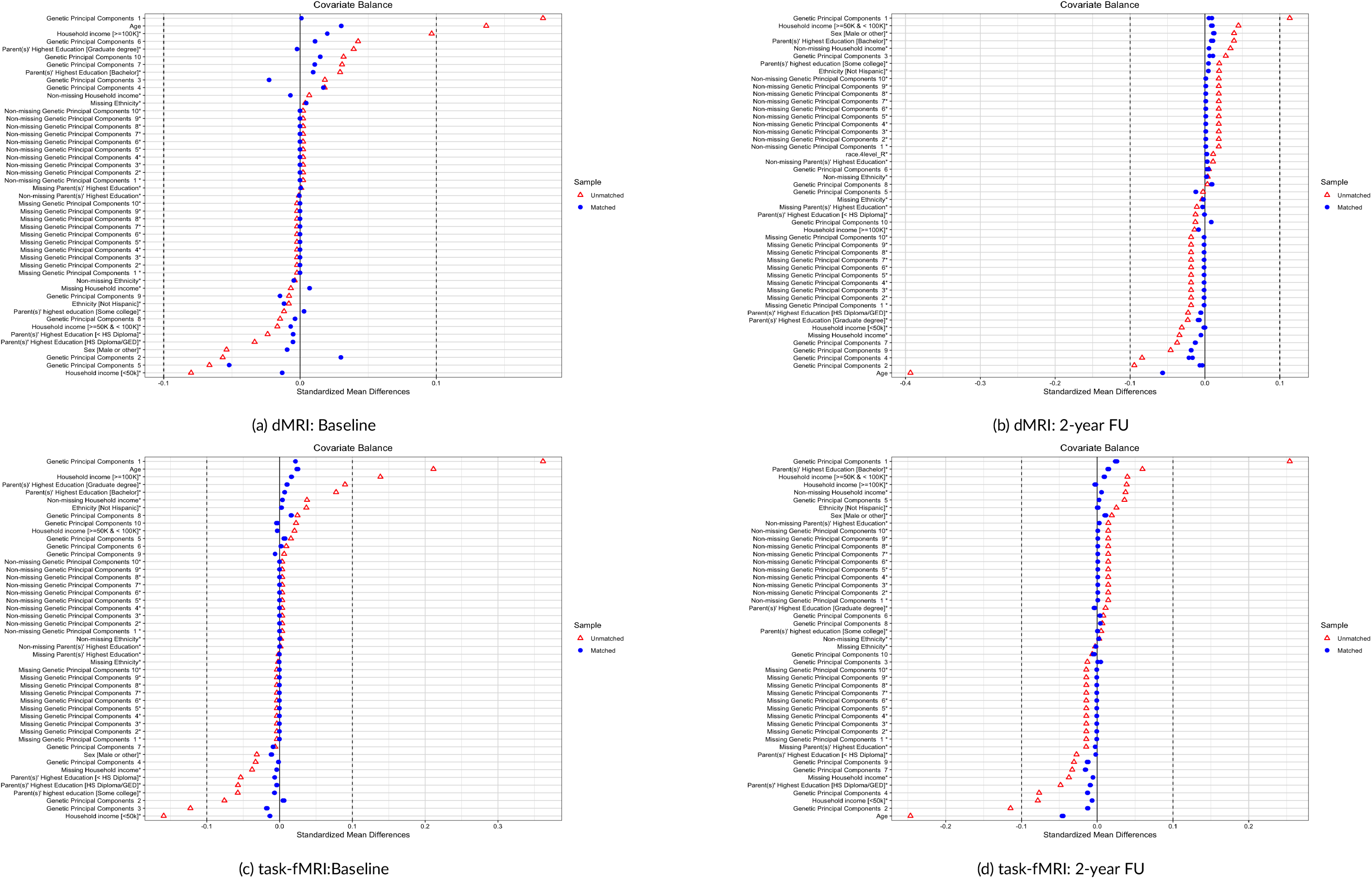
Covariates balance plots for Modified Model: dMRI and task-fMRI. Red triangles represent the unmatched sample and blue circles represent the matched sample. The closer the blue circles (matched sample) are to the vertical dashed line at zero, the better the balance between the treatment and control group for that covariate after matching

**FIGURE 2.**
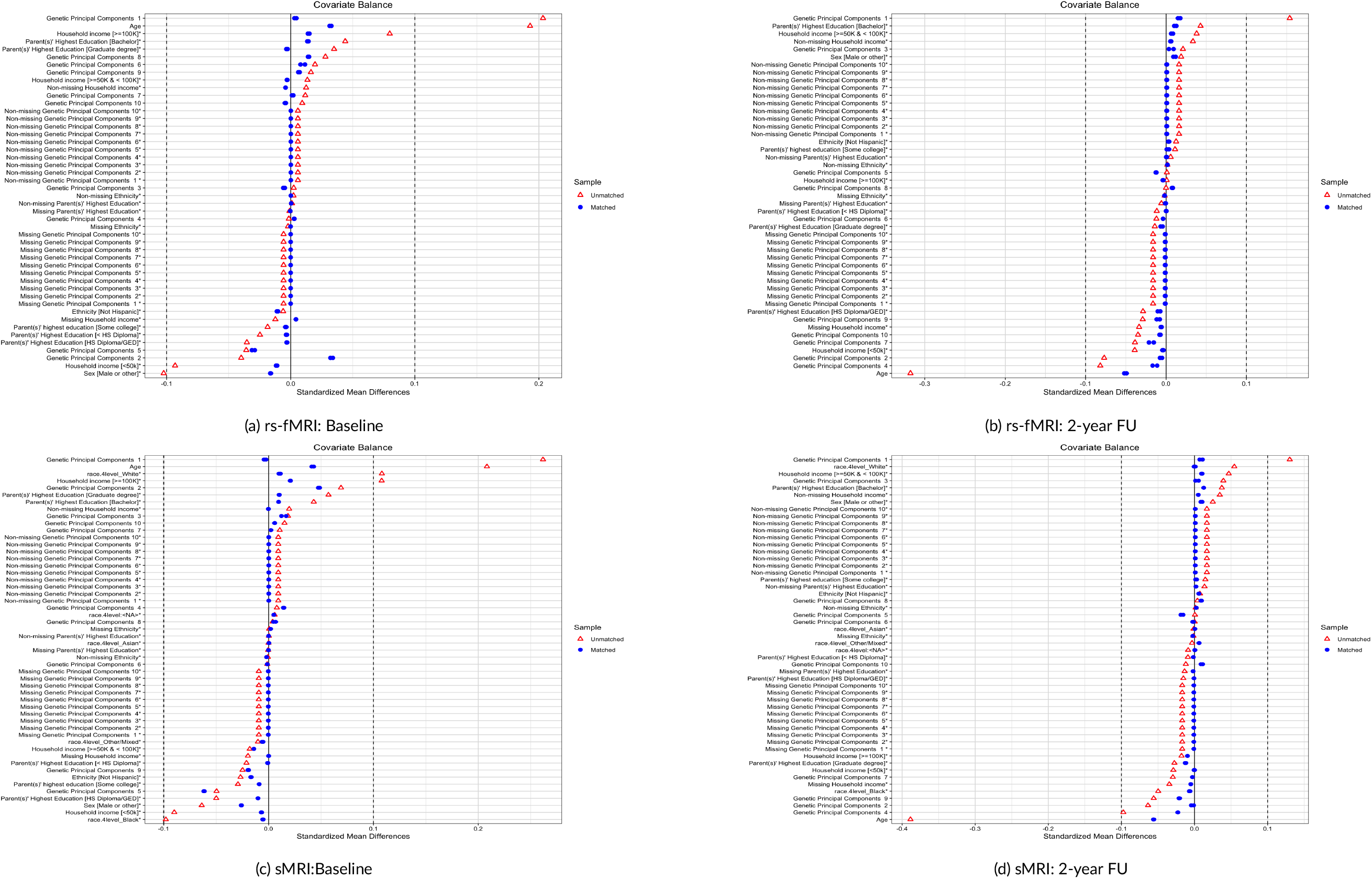
Covariates balance plots for Modified Model: rs-fMRI and sMRI. There are two sets of symbols: red triangles represent the unmatched sample and blue circles represent the matched sample. The closer the blue circles (matched sample) are to the vertical dashed line at zero, the better the balance between the treatment and control group for that covariate after matching

Figure 1 demonstrates the balance of dMRI and task-fMRI in baseline (left) and 2YRFU (right), respectively. The red triangles (unmatched sample) show the balance before matching, which often deviates more from zero, indicating less balance. The horizontal axis shows the standardized mean differences for each covariate, showing the differences in (standardized) means between the matched and unmatched groups, and the relative distance from the threshold of ±0.10[67]. Although there is debate regarding what value of a standardized difference denotes residual imbalance between treated and untreated subjects in the matched sample, research suggests that a threshold of 0.10 (or 10%) indicates a meaningful difference in covariate means between treatment and control group. The covariates are listed on the vertical axis, with the vertical dashed line at zero (0) representing perfect balance. The asterisks next the covariate names indicate that the mean differences of those variables have been standardized. As in Figure 1, Figure 2 shows how the covariates for rsfMRI and sMRI are balanced by the propensity score matching approach in baseline (left) and 2YR FU (right).

After evaluating propensity score estimates (PS) and their impact on data matching, we proceeded to apply propensity score weighting (PSW) as a missing data approach to model associations between age and longitudinal changes in brain metrics. For each MRI modality, propensity scores at baseline and at a 2-year follow-up (2YRFU) were transformed into inverse probability weights (calculated as the reciprocal of the propensity score 1/(1 − *P S*) and referred to as “PSW” for the remainder of these analyses).

For each participant in each modality, a final weight was constructed by multiplying the post-stratification weight from baseline with the PSWs from baseline and the 2YRFU. To mitigate the influence of extreme weights [68, 69], the final weight was trimmed by 1% on both sides [70]. These final weights were then included in the fitting of linear mixed-effects models. Equation 4 demonstrates this below:

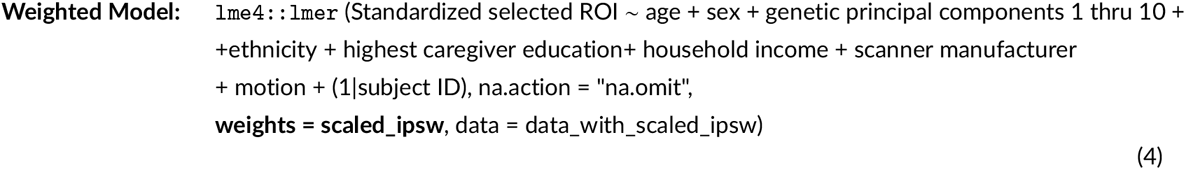

### 3.4 Comparison of Missing Data Approaches

### 3.5 Comparison of Point Estimates Across All Four Approaches

Figure 3 presents the point estimates and associated standard error bars for all four approaches for (a) Age, (b) Household income, and (c) Parental education, for all four modalities (dMRI, rs-fMRI, sMRI, and task-fMRI). Table 5 provides the same information in more precise terms for the age estimate, and also includes the intercept estimate. In general, the differences for the age estimates are negligible. Thus, for examining age effects using our Modified Model, the choice of missing data approach would not likely materially change the conclusions of the research question. However, this may not always be the case. For example, estimates of household income and parental education associations with longitudinal changes in brain metrics (Figure 3b and Figure 3c) are more variable across approaches. However, whether these are meaningful differences across approaches could only be determined with simulated data. Thus, we can’t make specific recommendations on which approach to use for particular research questions, but in the Discussion section we review the benefits and challenges of each missing data approach.

**FIGURE 3.**
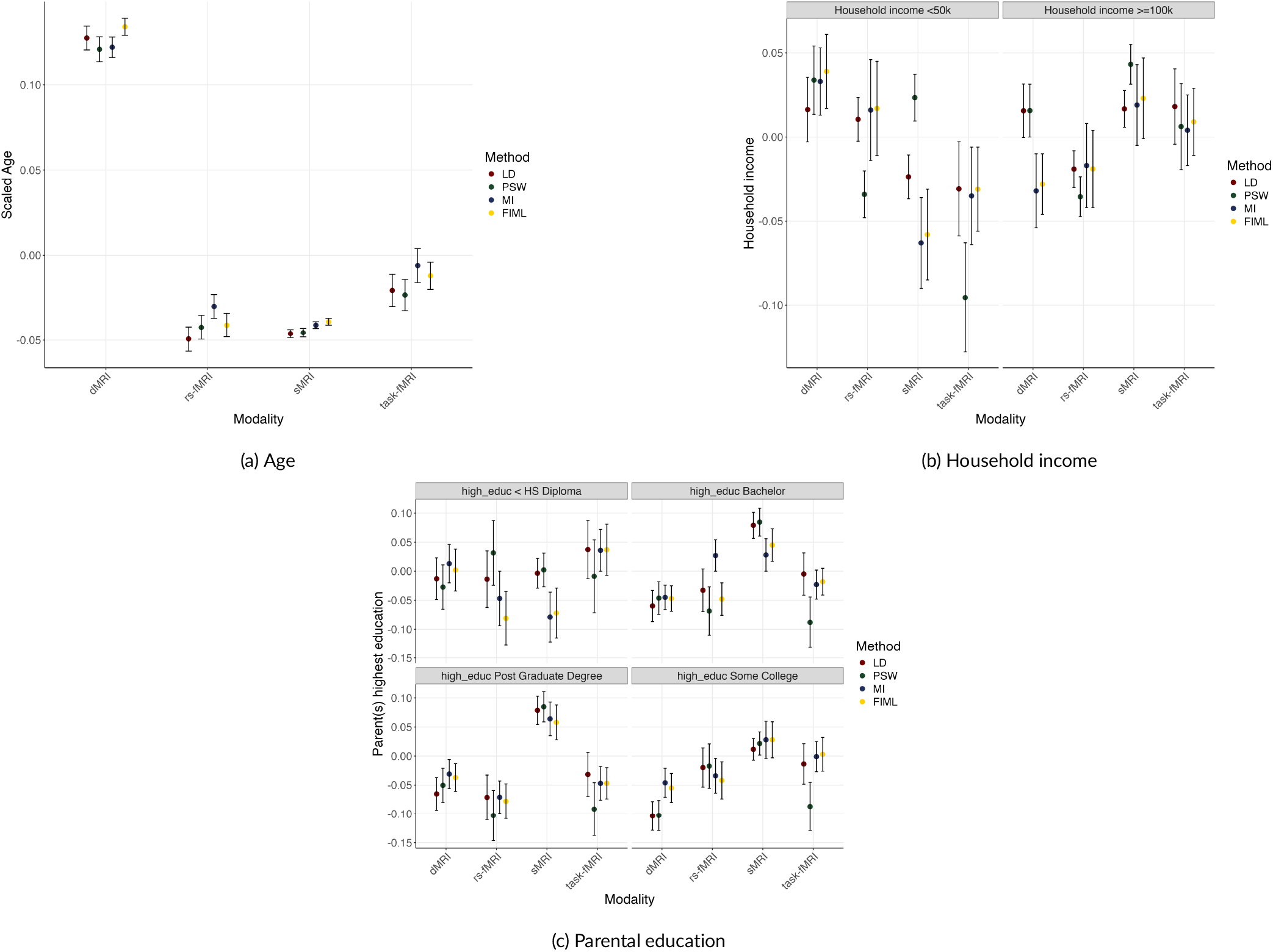
Point estimates and corresponding standard errors for the predictors of (a) age, (b) household income, and (c) highest education of parent(s), for each modality and estimating method. dMRI = Diffusion-weighted magnetic resonance imaging; task-fMRI = Task-related functional magnetic resonance imaging; rs-fMRI = Resting state magnetic resonance imaging; sMRI = Structural magnetic resonance imaging. LD = Listwise Deletion; PSW = Propensity Score Weighting. MI = Multiple Imputation. FIML = Full Information Maximum Likelihood.

**TABLE 5.** Comparison among Listwise Deletion, Propensity Score Weighting, Multiple Imputation, and Full Information Maximum Likelihood approaches for the Modified Model for each Outcome.

|  | dMRI |  |  |  | task-fMRI |  |  |  | rs-fMRI |  |  |  | sMRI |  |  |  |
| --- | --- | --- | --- | --- | --- | --- | --- | --- | --- | --- | --- | --- | --- | --- | --- | --- |
|  | LD | PSW | MI | FIML | LD | PSW | MI | FIML | LD | PSW | MI | FIML | LD | PSW | MI | FIML |
| Fixed Effect Intercept | 0.6776 *** | 0.6885 *** | -0.853*** | -0.786 *** | 0.0157 | 0.1245 * | 0.036 | 0.007 | 0.0367 | 0.0839 | 0.163*** | 0.155 *** | -0.2025*** | -0.2816*** | -0.189*** | -0.196 *** |
|  | (0.0351) | (0.0360) | (0.026) | (0.038) | (0.0405) | (0.0484) | (0.028) | (0.036) | (0.0408) | (0.0162) | (0.037) | (0.041) | (0.0285) | (0.0293) | (0.028) | (0.037) |
| Scaled Age | 0.1274 *** | 0.1208 *** | 0.122 *** | 0.134 *** | -0.0206 * | -0.0233 * | -0.006 | -0.012 | -0.0493 *** | -0.0423 *** | -0.030 *** | -0.041*** | -0.0461 *** | -0.0455 *** | -0.041 *** | -0.039 *** |
|  | (0.0070) | (0.0073) | (0.006) | (0.005) | (0.0095) | (0.0092) | (0.008) | (0.008) | (0.0072) | (0.0071) | (0.007) | (0.007) | (0.0024) | (0.0026) | (0.002) | (0.002) |

Note. LD = Listwise Deletion; PSW = Propensity Score Weighting; MI = Multiple Imputation; FIML = Full Information Maximum Likelihood. \* $p < 0.05$ ; \*\* $p < 0.01$ ; \*\*\* $p < 0.001$ (uncorrected).

## 4 DISCUSSION AND RECOMMENDATIONS FOR FUTURE WORK

In this paper, we explore how to apply three different missing data methods to the analysis of large longitudinal data sets, using the ABCD^®^ Study as an example. Thus, in addition to exploring these methods, we provide Supplemental Code so that researchers can use these methods in their own analysis. In our exploration, we found that all four methods, even listwise deletion, yielded similar results for the main predictor of interest, i.e., age. If this is the case, a researcher may question the value of using these additional methods, which are indeed more time-consuming and difficult to implement than the default listwise deletion. Yet one cannot know ahead of time whether listwise deletion introduces bias, or which missing data mechanisms may be at play. Indeed, it is possible that the large sample size buffers the effects of missingness for this particular predictor (age) or these particular outcomes. However, if a researcher were addressing research questions about a small subsample of the data (e.g., on the predictor side, asking questions about a rare disorder, or about exposure to rare risk factors for health outcomes, or on the outcome side, about rare behavioral or brain outcomes that might be zero-inflated), or if the degree of missingness is large, missingness may have more of an impact than we see here.

In fact, our results demonstrated that the impact of different missing data approaches on point estimates varied by predictor. Results with household income and parental education, which are some of the constructs with the most missing data in the ABCD^®^ Study [8], led to more variability in point estimate interpretations. Prior work in this sample [8] and in the broader survey methodology literature [71, 72] suggests that we might expect more variability by analytic approach for variables that are more closely tied to missingness and/or for which there are larger amounts of missing data. We hope these examples reiterate the importance of carefully considering the missing data mechanism(s) for each research question individually. That said, our analysis is not designed to compare across different missing data options. Such comparative studies exist already in the literature[40, 41, 50, 49] and require testing simulated data, which establish a “ground truth.” We don’t have access to the ground truth of these statistical models. Instead, a description of the challenges we encountered is likely most useful for researchers.

In general, for simple models, the available software are straightforward to use. But here, the longitudinal nature of the data coupled with the size of the dataset makes analyses of these large longitudinal data sets especially challenging. First, for ABCD^®^, as of the release we analyzed (5.1), the data are available in stacked or long format, and as a single data set pose computational challenges if considered as a whole. Thus, manipulation of the data format was necessary for some approaches to change the data from stacked/long to wide format. We provide code for this, which we hope will be a useful resource. Second, the complexity of the model determines the probability that the model will converge. Indeed, we encountered model convergence issues, but the challenges we encountered differed somewhat across approaches.

Making clear recommendations for which missing data approach researchers should implement is difficult because the appropriate method may differ depending on missing data patterns and the question of interest. Instead, we discuss pros and cons of the different approaches. For propensity score weighting, the main challenges lie in extreme weights that will increase variance estimates and result in overly conservative standard errors. Therefore, wave-specific weights need be examined for outliers, and may need to be trimmed before combining weights across waves of a longitudinal dataset. As outlined in the Methods section, we trimmed the final weight by truncating weights at the 1st and 99th percentiles [70]. And yet, as with so many statistical quandries (e.g., what is a “meaningful” effect size; goodness-of-fit indices), more conservative thresholds have been used elsewhere [73]. Benefits were also apparent in the use of propensity scores. Most clearly, propensity score construction and implementation in statistical modeling is relatively straightforward [51], owing to well-developed and maintained open-source R packages, like the twang [62] package used here. The ps() mnps() functions that are integrated within twang provide rigorous and robust algorithms to balance two (or three) groups on a set of a-priori covariates.

For multiple imputation, categorical predictors with a large number of levels were a challenge. We ended up collapsing these into a smaller number of variables (e.g., we collapsed scanner ID into a three-level scanner manufacturer variable). Other options are available here though. For example, variables with a number of levels such as scanner ID or site could be considered as random effects instead of fixed effects, if this makes sense in the analysis framework. In some cases, changing variables may change the operationalization of a construct. For example, genetic principal component scores replaced the common racial-ethnic identity categorical variable in this analysis. In ABCD^®^, several racial categories have been proposed [74], and in the past, 4-level and 6-level categorical variables have been part of the different ABCD^®^ data releases. But there is legitimate discussion about whether these operationalizations are ideal reflections of the construct of interest [75, 76]. At any rate, the choices about the initial variable for a certain construct, and any variables established to deal with convergence issues, must be determined in the context of the research question under investigation. Despite the challenges, multiple imputation has some unique benefits. For example, multiple imputation, as implemented in BLIMP, provides measures of model convergence. The PSR diagnostic indicates that the imputation procedure arrived at a consistent answer with respect to the imputed data across more than one Markov chain. This, coupled with the adjusted standard error, provides some sense of the repeatability of the imputation estimates. The FMI metric also enables evaluation of the degree to which missingness impacts the estimation of specific variables in the model, which is useful for understanding the model more generally.

FIML also comes with associated challenges and benefits, and again this depends somewhat on the model and data structure. Because it seems straightforward, researchers often ask why the default missing data approach is not to simply change the estimation procedure to FIML. At the outset, one barrier to this seemingly-simple strategy may be software. Because standard software for multilevel modeling (e.g., fitting linear mixed models in SPSS or using the lme or lmer functions in R) generally requires complete cases ^2^, individuals hoping to use these methods are forced to use either listwise deletion, propensity score weights applied to the complete cases, or multiple imputation to fill in all missing values. To use FIML estimation, one must switch to a software package capable of estimating multilevel structural equation models (MSEMs) with missing data, such as Mplus. Although univariate multilevel models are special cases of more general models and can readily be translated into an MSEM model, users unfamiliar with these models may face a learning curve in attempting to master different software interfaces, estimation defaults, and syntax language conventions (e.g., %within% and %between% specification in Mplus).

Nonetheless, if the model is simple, the data are cooperative, and the investigator is familiar with the software, FIML may be a viable and user-friendly approach. But this may not always be the case. One challenge when implementing FIML (in both single- and multilevel settings) is that all (exogenous) predictors in a given model must be brought into the multivariate normal likelihood function and treated as estimated quantities rather than fixed quantities as they would be in a regression-based analyses with complete data. That is, to use all available information on the X’s as well as the Y variable in an analysis, distributional assumptions (e.g., multivariate normality) must be imposed upon the X’s. When all predictors in a model are plausibly continuous, this poses no problem. But when dummy-coded predictors are included in a model (such as the sex, race-ethnicity, and scanner ID dummy variables in our example models), this creates issues. Not only are dichotomous variables inherently discrete and non-normal to a degree that is fundamentally at odds with an assumption of multivariate normality, each dummy variable’s variance (*p*\*(1 – *p*)) is a transformation of its mean (*p*, the proportion of 1’s on the dummy variable). For these reasons, bringing variables like these into the FIML likelihood function generally provokes program warning messages alerting the user to possible model non-identification, untrustworthy standard errors, and non-positive definiteness. Indeed, these messages occurred in every FIML model reported here. Reassuringly, simulations by Muthén et al. (2016) suggest that, despite their obvious departures from multivariate normality, FIML estimation with exogenous dummy variables may remain accurate nonetheless, implying that resulting warning messages may be safely ignored [77]. Yet, we believe that disregarding program error messages is generally a perilous strategy that should be undertaken with extreme caution, if not avoided altogether. Treating exogenous dummy variables as multivariate normally distributed variables is only one possible source of an error message. However, it is possible that multiple issues may be co-occurring in the syntax specification of any complex model. Habitually treating program warnings as safe to ignore risks failing to diagnose additional issues that may be present. Thus, we cannot recommend this strategy without serious reservation. However, if these challenges are not present (for example, if all predictors are ordinal or continuous and do not provoke such error messages), a main benefit of FIML is that the procedure is effective at dealing with missing data while also being simple to describe, staying close to the description one would use to explain a standard multilevel model.

In summary, the missing data approaches we showed establish different challenges and benefits, and we hope the examples, recommendations for general best practices, and example code we provide help researchers examining large longitudinal data sets. It is encouraging, from one perspective, that the different approaches yielded similar results for the focal predictor of age (and, to some extent, socioeconomic constructs) because it suggests they converge on a consistent solution. This was the case here, but it may not always be the case, and it is difficult to guess when results from analyses with missing data will be impartial or instead subject to substantial bias. Regardless, simulation studies, which are able to determine a “ground truth”, establish the effectiveness of these approaches. Now investigators can use this resource in the examination of their own research questions in the context of large longitudinal data sets like the ABCD^®^ study.

## Acknowledgements

Thank you to Craig Enders who answered some questions about model convergence in BLIMP. Thank you also to Brian Kim who answered some questions with regards to combining propensity scores across timepoints. Data used in the preparation of this article were obtained from the Adolescent Brain Cognitive Development (ABCD) Study (https://abcdstudy.org), held in the NIMH Data Archive (NDA). This is a multisite, longitudinal study designed to recruit more than 10,000 children age 9-10 and follow them over 10 years into early adulthood. The ABCD Study is supported by the National Institutes of Health and additional federal partners under award numbers U01DA041048, U01DA050989, U01DA051016, U01DA041022, U01DA051018, U01DA051037, U01DA050987, U01DA041174, U01DA041106, U01DA041117, U01DA041028, U01DA041134, U01DA050988, U01DA051039, U01DA041156, U01DA041025, U01DA041120, U01DA051038, U01DA041148, U01DA041093, U01DA041089, U24DA041123, U24DA041147. A full list of supporters is available at https://abcdstudy.org/federal-partners.html. A listing of participating sites and a complete listing of the study investigators can be found at https://abcdstudy.org/consortium_members/. ABCD consortium investigators designed and implemented the study and/or provided data but did not necessarily participate in the analysis or writing of this report. This manuscript reflects the views of the authors and may not reflect the opinions or views of the NIH or ABCD consortium investigators. The ABCD data repository grows and changes over time. The ABCD data (release 5.1) used in this report came from DOI: http://dx.doi.org/10.15154/z563-zd24.

## Conflict of Interest

The authors report no conflicts of interest.

## Code Availability

All code for this study is available at https://osf.io/q8v5s/.

## Footnotes

1 Note the mnps function in the twang package (v2.5) [62] is an extension that allows for balancing across three groups

2 Note that multilevel models do allow for unbalanced cluster sizes. As a result, these models readily accommodate situations in which in-dividuals (clusters) are missing data on certain time points (level-1 observations) in a longitudinal analysis. In this context, individuals with missing time points will have fewer rows of data than individuals with complete data at all timepoints. But all data that they do have must be complete. Any rows with explicit missing values (e.g., NA values in R) will be listwise deleted from the analysis.

## Notes

https://osf.io/q8v5s/

### Competing Interest Statement

The authors have declared no competing interest.

