## Supplemental Materials for "Missing data approaches for longitudinal neuroimaging research: Examples from the Adolescent Brain and Cognitive Development (ABCD) Study"

### | Missingness patterns of sociedemographical variables

**TABLE S1** Patterns of Missingness by sociedemographics

| Family Income (Longitudinal) | Parents Highest Education (Longitudinal) | Race (4-level, Baseline only) | Ethnicity (Hispanic, Baseline only) | Baseline | 2 Year Follow-up |
|---|---|---|---|---|---|
| Missing | | | | 1017 | 964 |
| | | Missing | | 438 | 396 |
| | | | Missing | 153 | 132 |
| | Missing | | | 17 | 1776 |
| Missing | Missing | | | 11 | 255 |
| Missing | | Missing | | 72 | 71 |
| Missing | | | Missing | 26 | 20 |
| | Missing | Missing | | 1 | 49 |
| | Missing | | Missing | 1 | 16 |
| | | Missing | Missing | 19 | 16 |
| Missing | Missing | Missing | | 0 | 13 |
| Missing | Missing | | Missing | 1 | 7 |
| Missing | | Missing | Missing | 2 | 7 |
| | Missing | Missing | Missing | 0 | 5 |
| Missing | Missing | Missing | Missing | 0 | 3 |

The table above documents the number of missing data points across different demographic variables and time points in a study. Under each demographic category, the table shows the number of missing data points at baselineand at 2-year follow up. For instance, there were 1014 missing data points for the Family Income at baseline and 898 at the 2-year follow up. Notably, there's a high number of missing data points for ethnicity (Hispanic) at baseline (1711), which has substantially improved at the 2-year follow up (131). The table is used to assess the completeness of the data collected in the study.

### | Missing data mechanisms of rs-fMRI and sMRI

For fMRI data from the resting state at baseline, age (OR = 1.35, $p < 0.001$), sex (Female, OR = 1.52, $p < 0.001$; ref = Male or intersex), race (Asian, OR = 0.50, p = 0.035; Black, OR =0.64 $p < 0.001$; ref = White), annual household income (<50k, OR = 0.79, p = 0.005; ref = [>=100k]), and caregiver highest education (< HS Diploma, OR = 0.68, p = 0.002; HS Diploma/GED, OR = 0.73, p = 0.001; ref = Bachelor), were significantly associated with missingness. The missing resting-state fMRI data at 2 years of follow-up was associated with age (OR = 0.61 $p < 0.001$; ref = Male or intersex), sex (Female, OR =0.90, p = 0.023; ref = Male or intersex), race (Black, OR = 0.58, $p < 0.001$ ; Other/Mixed, OR = 0.88, p = 0.048; ref = White), ethnicity (Hispanic, OR = 0.87, p = 0.036; ref = non-hispanic), annual household income ($\geq$ 50k&<100k, OR = 1.19, p = 0.004; ref = [>=100k]), and caregiver highest education (HS Diploma/GED, OR = 0.80 p = 0.012; Post Graduate degree, OR = 0.85, p = 0.004; ref = Bachelor).

sMRI evinced similar patterns of associations between sociodemographic constructs and missingness in both waves. For example, age (OR =1.40, $p < 0.001$), sex (Female, OR = 1.29, $p < 0.001$; ref = Male or intersex), race (Black, OR

= 0.61, p < 0.001 ; Other/Mixed, OR = 0.78, p = 0.010; ref = White), ethnicity (Hispanic, OR = 1.35, p = 0.002; ref = non-hispanic), annual household income (<50k, OR = 0.79, p = 0.030;≥ 50k&<100k, OR = 0.78, p = 0.005; ref = [>=100k]), and caregiver highest education (< HS Diploma, OR = 0.70, p = 0.033; HS Diploma/GED, OR = 0.71, p = 0.009; ref = Bachelor), were significantly associated with missing sMRI data at baseline. At the 2 year follow-up, age (OR = 0.54, p < 0.001), sex (Female, OR = 0.88 p = 0.008; ref = Male or intersex), race (Black, OR = 0.61, p < 0.001; ref = White) , ethnicity (Hispanic, OR = 0.84, p = 0.015; ref = non-hispanic), annual household income ($\geq 50k\&< 100$k, OR = 1.26, p < 0.001; ref = [>=100k]), and caregiver highest education (Post graduate degree, OR = 0.82, p = 0.001; ref = Bachelor) were significantly associated with missing DWI data.
